# Evaluating The Efficacy Of Endotracheal Epinephrine Administration At Standard Versus High Dose During Resuscitation Of Severely Asphyxiated Newborn Lambs: A Randomized Preclinical Study

**DOI:** 10.1101/2023.02.28.530542

**Authors:** Graeme R. Polglase, Yoveena Brian, Darcy Tantanis, Douglas A. Blank, Shiraz Badurdeen, Kelly J. Crossley, Martin Kluckow, Andrew W. Gill, Emily Camm, Robert Galinsky, Nils Thomas Songstad, Claus Klingenberg, Stuart B. Hooper, Calum T. Roberts

**Affiliations:** The Ritchie Centre, Hudson Institute of Medical Research and Department of Obstetrics and Gynaecology, Monash University, Melbourne, VIC, Australia; Department of Paediatrics, Monash University, Melbourne, VIC, Australia; Monash Newborn, Monash Children’s Hospital, Melbourne, VIC, Australia; Newborn Research Centre, The Royal Women’s Hospital, Melbourne, Australia; Department of Neonatology, Royal North Shore Hospital & University of Sydney, Sydney, NSW, Australia; Centre for Neonatal Research and Education, The University of Western Australia, Subiaco, WA, Australia; University Hospital of North Norway, Tromsø, Norway

**Keywords:** Resuscitation, Infant, newborn, Animal, Epinephrine, Infusions

## Abstract

**Background:** Epinephrine treatment is recommended during neonatal resuscitation, if ventilation and chest compressions are ineffective. Endotracheal administration is an option, if the preferred intravenous route is unavailable. We aimed to determine the efficacy of endotracheal epinephrine for achieving return of spontaneous circulation (ROSC), and maintaining physiological stability after ROSC, at standard and higher dose, in severely asphyxiated newborn lambs.

**Methods:** Near-term fetal lambs were instrumented for physiological monitoring, and asphyxiated until asystole. Resuscitation was commenced with ventilation and chest compressions as per ILCOR recommendations. Lambs were randomly allocated to: IV Saline placebo (5 ml/kg, n=6), IV Epinephrine (20 micrograms/kg, n=9), Standard-dose ET Epinephrine (100 micrograms/kg, n=9), and High-dose ET Epinephrine (1 mg/kg, n=9). After three allocated treatment doses, rescue IV Epinephrine was administered if ROSC had not occurred. Lambs achieving ROSC were ventilated and monitored for 60 minutes before euthanasia. Brain histology was assessed for micro-hemorrhage.

**Results:** ROSC in response to allocated treatment (without rescue IV Epinephrine) occurred in 1/6 Saline, 9/9 IV Epinephrine, 0/9 Standard-dose ET Epinephrine, and 7/9 High-dose ET Epinephrine lambs respectively. Three Saline, six Standard-dose ET Epinephrine, and one High-dose ET Epinephrine lambs achieved ROSC after rescue IV Epinephrine. Blood pressure during CPR increased after treatment with IV Epinephrine and High-dose ET Epinephrine, but not Saline or Standard-dose ET Epinephrine.

After ROSC, both ET Epinephrine groups had lower pH, higher lactate, and higher blood pressure than the IV Epinephrine group. Cortex micro-hemorrhage was more frequent in the High-dose ET Epinephrine lambs (8/8 lambs examined, versus 3/8 in IV Epinephrine lambs).

**Conclusions:** The currently recommended dose of ET Epinephrine was ineffective in achieving ROSC. In the absence of convincing clinical or preclinical evidence of efficacy, use of ET Epinephrine at this dose may not be appropriate.

High-dose ET Epinephrine requires further evaluation before clinical translation.

## Introduction

Globally, birth asphyxia secondary to intrapartum-related conditions is a major cause of neonatal mortality (second only to prematurity), resulting in more than 700,000 deaths annually.(1) Amongst survivors, >400,000 infants per year will be neurologically impaired as a result of birth asphyxia.(2) Infants exposed to asphyxia are likely to require resuscitation at birth. Although the majority of infants that receive resuscitation will respond to basic measures such as provision of positive pressure ventilation, those more severely affected by asphyxia will require advanced measures. Current recommendations advise that cardiopulmonary resuscitation (CPR) with chest compressions is commenced for infants who are asystolic, or bradycardic with heart rate <60 bpm, despite adequate ventilation. For those most severely compromised, the delivery of epinephrine is often essential to achieve return of spontaneous circulation (ROSC). Although required in only a minority, those infants treated with epinephrine have particularly high rates of neurodevelopmental impairment and death.(3)

Neonatal resuscitation guidelines include several potential routes for epinephrine administration, with intravenous (IV) administration via an umbilical venous catheter (UVC) preferred.(4, 5) However, gaining IV access may be time-consuming, or challenging to achieve with limited numbers of resuscitation staff, thus delaying epinephrine administration and placing the newborn at increased risk of prolonged hypoxia and ischemia due to poor cardiac function. Alternatively, epinephrine can be administered via an endotracheal tube, which may be placed early during the resuscitation for airway management. (4, 5) In simulated neonatal resuscitation, endotracheal intubation is achieved four minutes earlier than UVC placement, representing a clear advantage in achieving timely administration of epinephrine.(6)

However, epinephrine can only manifest its desired cardiovascular effects if it is sufficiently available within the systemic circulation. Early preclinical data suggested that endotracheal epinephrine administration could effectively achieve ROSC.(7) Studies in anaesthetised dogs showed that endotracheally administered epinephrine was effectively absorbed into the plasma,(8) but studies in both animals with spontaneous circulation, and those receiving CPR after cardiac arrest, suggested that increased dosing was required in comparison with the recommended IV dose to produce an elevation in plasma levels.(9, 10) These studies were all conducted in older animals rather than newborns, so did not allow for important physiological differences present in the newborn infant immediately after birth. Prior to lung aeration, newborn infants have low pulmonary blood flow with a patent ductus arteriosus, and their airways are filled with liquid.(11, 12) These differences could potentially have significant effects on absorption and bioavailability of any drug administered into the trachea, including epinephrine. More recent preclinical work in newborn lambs has shown that administration of endotracheal epinephrine at the current recommended maximum dose (100 micrograms/kg) results in lower plasma levels and lower rates of ROSC without the use of ‘rescue’ IV epinephrine, when compared with standard dose IV epinephrine (10-30 micrograms/kg).(13, 14)

Clinical outcome data for the use of endotracheal epinephrine are limited: a systematic review conducted in 2020 identified four observational studies that were suitable for inclusion, with no randomised trials conducted in neonates.(15) The included observational studies contained 94 infants treated with endotracheal epinephrine. In the two largest studies combined, 54/74 infants (73%) receiving endotracheal epinephrine did not respond, and subsequently received at least one IV epinephrine dose.(16, 17)

Resuscitation guidelines must take several factors into consideration, including ease of use, time for device insertion, and effectiveness of treatment, when recommending routes of epinephrine administration in the clinical setting (Figure 1). Current data suggest that the endotracheal route may provide some advantage in speed of administration, but has relatively poor efficacy. It is conceivable that the dose of endotracheal epinephrine required to achieve ROSC in the newborn infant is substantially higher than that currently recommended, and that this may explain the reduced rates of ROSC seen in both preclinical and clinical reports. However, endotracheal epinephrine administration appears to produce a sustained and amplified peak plasma level after spontaneous circulation has been restored,(13, 14) so such an escalated dose may result in adverse effects in the post-resuscitation period.

**Figure 1.**
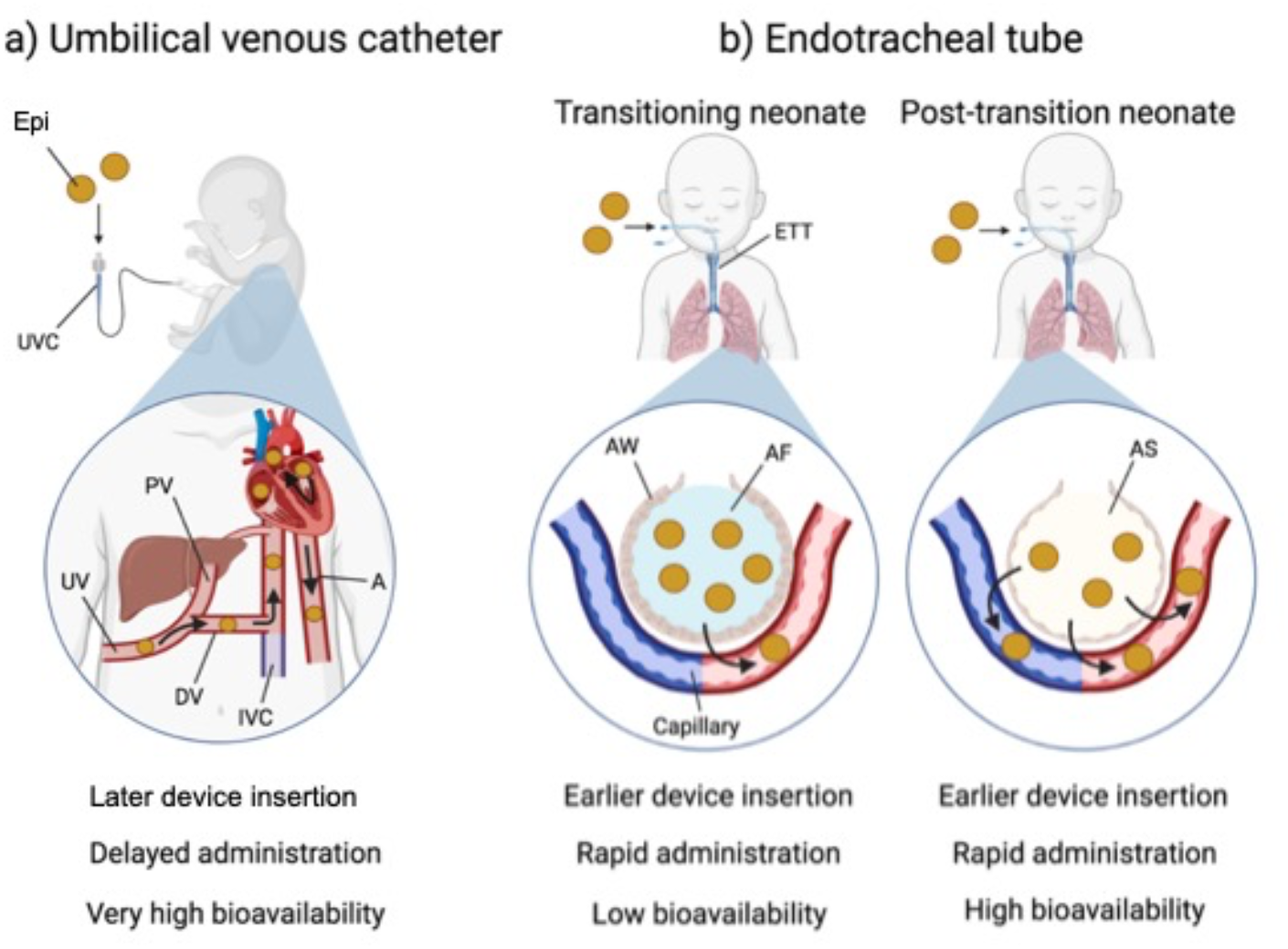
Contrasting features of intravenous and endotracheal epinephrine administration in neonatal resuscitation. a) IV epinephrine administration by umbilical venous catheter results in very high bioavailability in the central circulation, but may be more challenging or time-consuming. b) Endotracheal epinephrine administration may be achieved earlier in resuscitation but with lower resulting bioavailability. Epi: Epinephrine; UVC: Umbilical venous catheter; UV: Umbilical vein; PV: Portal vein; DV: Ductus venosus; IVC: Inferior vena cava; A: Aorta; ETT: Endotracheal tube; AW: Alveolar wall; AF: Alveolar fluid; AS: Alveolar space.

We therefore aimed to determine the efficacy of endotracheal epinephrine for restoring cardiac function, and maintaining physiological stability after ROSC, at both a standard dose and a higher dose, in severely asphyxiated newborn lambs. We hypothesised that regardless of dose, the endotracheal route would be less effective than the IV route in achieving ROSC.

## Methods

Monash Medical Centre Animal Ethics Committee A, Monash University (MMCA2020/04) approved all experimental procedures. The study was conducted in accordance with the National Health and Medical Research Council of Australia’s guidelines, and ARRIVE guidelines(18).

### Instrumentation and delivery

Pregnant Border-Leicester ewes (*Ovis Aries*) at (139 ± 2d) days gestation (mean±SD; term ∼148 days) were initially anaesthetised with intravenous thiopentone sodium (20 mg/kg) and anaesthesia was maintained, following intubation, by inhalation of isofluorane (1.5-5%) in air/oxygen; the gas mixture was adjusted to maintain maternal arterial oxygen saturations (SaO_2_) of >95%.

The fetus was partially exteriorised (head, chest and forelimbs) from the uterus and flow probes of appropriate size were placed around the left main pulmonary artery and carotid artery (Transonic Systems, Ithaca, NY, USA). Heparinized saline-filled catheters were inserted into a carotid artery, brachial artery and jugular vein as described previously.(19) The carotid artery catheter was connected to a pressure transducer (PD10; DTX Plus Transducer, Becton Dickinson, Singapore) to measure systemic blood pressure. The lamb was completely removed from the uterus and a rectal temperature probe was placed. A peripheral oxygen saturation (SpO_2_) probe (Masimo, Radical 4, CA, USA) was placed around the right forelimb and a Near Infrared Spectroscopy (NIRS) Optode (Casmed Foresight, CAS Medical Systems Inc, Branford, CT, USA) was placed over the scalp to continuously measure cerebral tissue oxygen saturation (SctO_2_). Blood pressures and flows, temperature and oxygen saturation were digitally recorded throughout the study (1kHz, Powerlab; ADInstruments, Castle Hill, NSW, Australia). The fetal trachea was intubated with a 4.5mm cuffed endotracheal tube, lung liquid was drained passively and the endotracheal tube clamped.

The umbilical cord was clamped and the lamb was weighed before it was transferred to an infant warmer (Fisher and Paykel Healthcare, Auckland, New Zealand). Asphyxia of the lamb was induced by withholding respiratory support and progressed until the mean blood pressure was reduced to ∼0 mmHg, and no discernible activity was visible on the blood pressure/flow traces, as per previous studies.(19)

### Treatment Allocation

Immediately prior to surgery, lambs were randomly allocated, using a web-based random sequence generator (www.random.org/lists), to one of four treatment groups:

1. ‘Saline’ (n=6), treated with intravenous 0.9% saline placebo (5ml)
2. ‘IV Epinephrine’ (n=9), treated with intravenous epinephrine (20 micrograms/kg) according to standard neonatal resuscitation guidelines, followed by 0.9% saline flush (5ml)
3. ‘Standard-dose ET Epinephrine’ (n=9), treated with endotracheal epinephrine (100 micrograms/kg) at the maximum dose included in current neonatal resuscitation guidelines
4. ‘High-dose ET Epinephrine’ (n=9), treated with endotracheal epinephrine (1 mg/kg) at a dose 10 times greater than the maximum dose in current neonatal resuscitation guidelines

The researcher leading the resuscitation team was aware of the route of epinephrine administration, but blinded to the dose of ET epinephrine, which was provided as an identical volume (1ml/kg) of either 1:10,000 (standard dose) or 1:1,000 (High-dose) epinephrine solution.

### Resuscitation and Post-Resuscitation Care

Resuscitation was initiated in air using positive pressure ventilation via a T-piece device (Neopuff; Fisher and Paykel Healthcare, Auckland, New Zealand) with peak inflation pressure 30 cmH_2_O and end-expiratory pressure 5 cmH_2_O, targeting 60 inflations per minute. One minute after ventilation onset, chest compressions were initiated with a target of 90 compressions and 30 inflations per minute, and the fraction of inspired oxygen was increased to 1.00 as per resuscitation guidelines.(5).

One minute after initiation of CPR and every three minutes thereafter, saline or epinephrine was administered via the allocated route. ROSC was defined as diastolic blood pressure >20 mmHg and spontaneous heart rate at >100 bpm, and determined by the researcher leading the resuscitation. If ROSC was not achieved after three allocated treatment doses, two ‘rescue’ doses of standard dose IV Epinephrine could be administered. CPR ceased at 15 minutes in lambs that failed to achieve ROSC by this time. If ROSC was achieved, CPR and treatment administration ceased, and the lambs were ventilated (Dräeger Babylog 8000+, Dräeger, Lübeck, Germany) with heated humidified gas (F&P 950 System, Fisher and Paykel, Auckland, New Zealand) for a further 60 minutes.

In the two ET epinephrine groups, plasma samples were collected from the brachial artery before cord clamping (fetal), at end asphyxia, and at 3, 6, 9, and 15 min after ROSC, and epinephrine concentrations were determined by enzyme immunoassay (Epinephrine 162 Research ELISA, catalogue #BA E-5100; LDN, Germany).

Blood gas samples (ABL30, Radiometer, Copenhagen, Denmark) were collected every 3 minutes until 15 minutes, then at 20, 25, 30, 40, 50, and 60 minutes. During the first 10 minutes after ROSC, lambs received pressure-limited ventilation at 30/5 cmH_2_O, after which volume guarantee ventilation was commenced at tidal volume of 7ml/kg. Ventilation parameters were also digitally recorded. Ventilation settings were adjusted to target SpO_2_ 90-95% and PaCO_2_ 35-45 mmHg. Lambs received sedation to ensure comfort (Alfaxan 5-15mg/kg/hr in 5% dextrose).

Following the experiment, ewes and lambs were euthanized using an intravenous overdose injection of sodium pentobarbitone (100 mg/kg IV, Lethobarb, Virbac, Australia).

### Histological analysis of microbleeds

The brains from all lambs were removed post mortem, weighed and the two hemispheres separated along the midline. The cerebellum was removed from the brain at the level of the cerebellar peduncles. The whole brain was immersion-fixed in 10% formalin for ∼5 days at 4°C.

After fixation, the right hemisphere was cut into coronal slices (5mm in thickness), processed and wax embedded, then sectioned at the level of the parietal lobe (10-micron sections). Sections were mounted on Superfrost microscope slides for histological staining with a 0.5% Cresyl Violet and 1% Acid Fuchsin solution.

### Quantitative Analysis

Quantitative analysis was performed on coded slides (observer blinded to group) using image analysis software (ImageScope, Aperio technologies, Vista, CA, USA). Micro-bleeds were identified according to the following criteria:

1. Indistinguishable basement membrane
2. Random dispersion of erythrocytes in tissue
3. Irregular shape of perimeter

These criteria are characteristic of extravasation.(20) The number of microbleeds per slide were measured within four brain regions: cortex, white matter, deep grey matter (including thalamus) and hippocampus. All analysis was performed in two sections per animal.

### Statistical analysis

All lambs were analysed for the assessments made during CPR while only lambs achieving ROSC were included in the post-ROSC analysis. All baseline fetal and physiological data during CPR were compared using a one-way ANOVA (GraphPad Prism; GraphPad Software, CA, USA). A Fisher’s exact test was used to compare dichotomous outcomes. A one-way ANOVA was also used to analyse the number of microbleeds in different regions of the brain. A two-way repeated measures ANOVA with “group” and “time” as the factors was used to compare groups during CPR. A two-way repeated measures ANOVA with Holm-Sidak *post hoc* comparison was used to compare the blood gases, post-ROSC physiological data and plasma epinephrine concentration during the experiment. Statistical significance was accepted at p<0.05. Data are reported as mean ± standard deviation unless otherwise stated.

Eight animals per group were required to demonstrate a reduction in rate of ROSC from 100%, as expected with the IV Epinephrine group, to 50% in a comparator group, with 80% power and alpha = 5%. We planned, *a priori*, to include 9 animals per group to optimise availability of post-ROSC physiological data for analysis, assuming reduced survival in some treatment groups. Allocation to the Saline Group was ceased at six animals as, based on ROSC rates in this and a previous study,(14) this treatment was clearly ineffective and the investigators felt it was unethical to continue allocating animals to this group.

## Results

### FETAL CHARACTERISTICS

Fetal characteristics for each treatment group, and post-mortem organ weights used in blood flow analysis, are shown in Table 1. Blood gas parameters were similar in the four groups prior to initiation of the study.

**Table 1:**
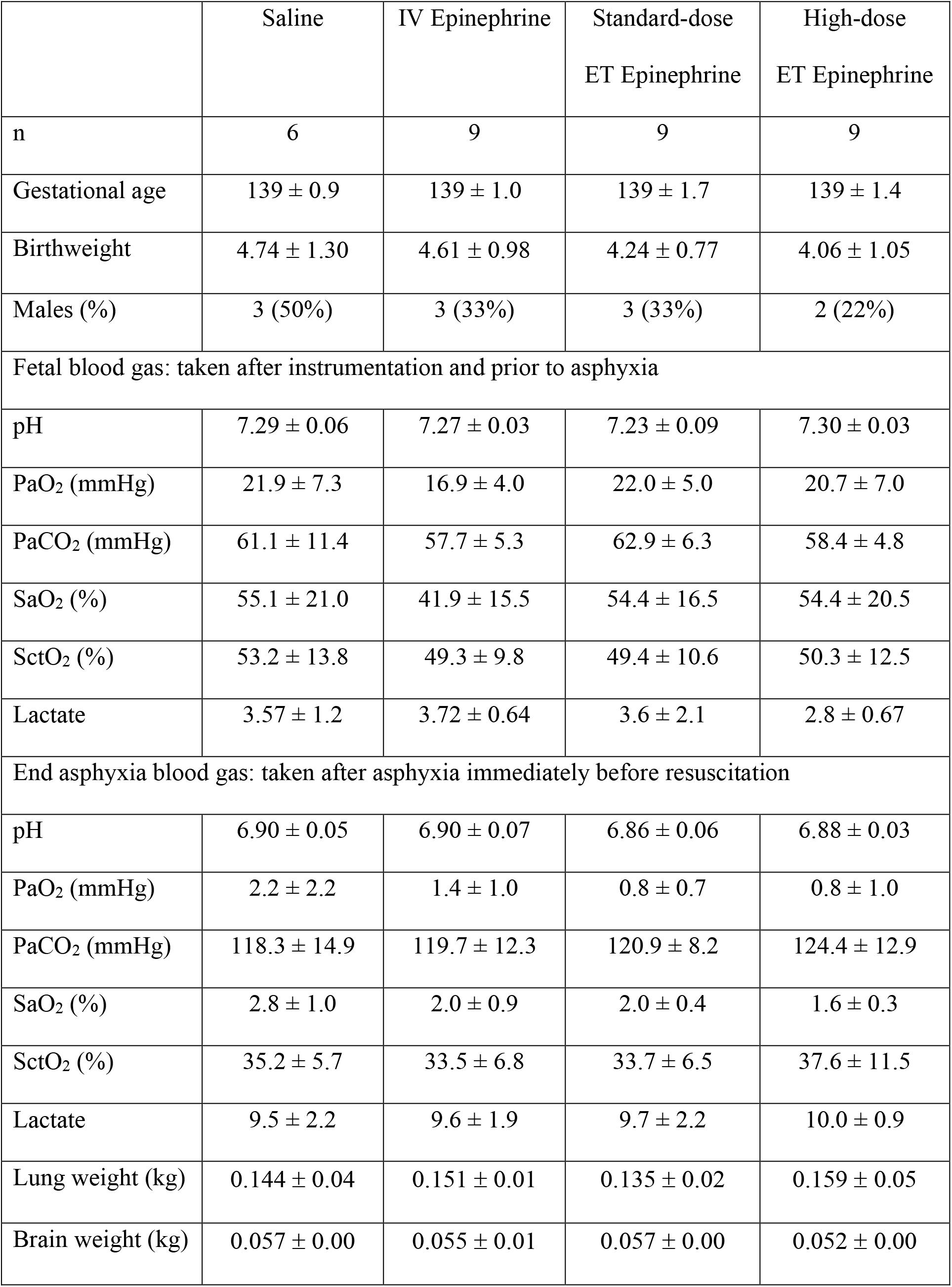
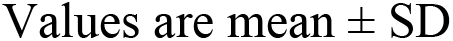
Fetal characteristics and blood gases, and post-mortem organ weights.

|  | Saline | IV Epinephrine | Standard-dose<br>ET Epinephrine | High-dose<br>ET Epinephrine |
| --- | --- | --- | --- | --- |
| n | 6 | 9 | 9 | 9 |
| Gestational age | 139 ± 0.9 | 139 ± 1.0 | 139 ± 1.7 | 139 ± 1.4 |
| Birthweight | 4.74 ± 1.30 | 4.61 ± 0.98 | 4.24 ± 0.77 | 4.06 ± 1.05 |
| Males (%) | 3 (50%) | 3 (33%) | 3 (33%) | 2 (22%) |
| Fetal blood gas: taken after instrumentation and prior to asphyxia |  |  |  |  |
| pH | 7.29 ± 0.06 | 7.27 ± 0.03 | 7.23 ± 0.09 | 7.30 ± 0.03 |
| PaO<sub>2</sub> (mmHg) | 21.9 ± 7.3 | 16.9 ± 4.0 | 22.0 ± 5.0 | 20.7 ± 7.0 |
| PaCO<sub>2</sub> (mmHg) | 61.1 ± 11.4 | 57.7 ± 5.3 | 62.9 ± 6.3 | 58.4 ± 4.8 |
| SaO<sub>2</sub> (%) | 55.1 ± 21.0 | 41.9 ± 15.5 | 54.4 ± 16.5 | 54.4 ± 20.5 |
| SctO<sub>2</sub> (%) | 53.2 ± 13.8 | 49.3 ± 9.8 | 49.4 ± 10.6 | 50.3 ± 12.5 |
| Lactate | 3.57 ± 1.2 | 3.72 ± 0.64 | 3.6 ± 2.1 | 2.8 ± 0.67 |
| End asphyxia blood gas: taken after asphyxia immediately before resuscitation |  |  |  |  |
| pH | 6.90 ± 0.05 | 6.90 ± 0.07 | 6.86 ± 0.06 | 6.88 ± 0.03 |
| PaO<sub>2</sub> (mmHg) | 2.2 ± 2.2 | 1.4 ± 1.0 | 0.8 ± 0.7 | 0.8 ± 1.0 |
| PaCO<sub>2</sub> (mmHg) | 118.3 ± 14.9 | 119.7 ± 12.3 | 120.9 ± 8.2 | 124.4 ± 12.9 |
| SaO<sub>2</sub> (%) | 2.8 ± 1.0 | 2.0 ± 0.9 | 2.0 ± 0.4 | 1.6 ± 0.3 |
| SctO<sub>2</sub> (%) | 35.2 ± 5.7 | 33.5 ± 6.8 | 33.7 ± 6.5 | 37.6 ± 11.5 |
| Lactate | 9.5 ± 2.2 | 9.6 ± 1.9 | 9.7 ± 2.2 | 10.0 ± 0.9 |
| Lung weight (kg) | 0.144 ± 0.04 | 0.151 ± 0.01 | 0.135 ± 0.02 | 0.159 ± 0.05 |
| Brain weight (kg) | 0.057 ± 0.00 | 0.055 ± 0.01 | 0.057 ± 0.00 | 0.052 ± 0.00 |

Values are mean ± SD

### RESPONSE TO TREATMENT AND SURVIVAL

Rates of ROSC, in response to allocated treatment alone, and following IV ‘rescue’ Epinephrine use if needed, are shown in Table 2. While overall survival did not differ significantly, higher rates of ROSC in response to allocated treatment were achieved in the IV Epinephrine and High-dose ET Epinephrine groups. Three lambs that achieved ROSC after rescue IV Epinephrine in the Standard-dose ET Epinephrine group subsequently had cardiac arrest prior to study completion, with 3/9 surviving to study end at 60 min.

**Table 2.** Response to treatment, survival, and brain histology.

|  | Saline (n=6) | IV<br>Epinephrine<br>(n=9) | Standard dose<br>ET Epinephrine<br>(n=9) | High-dose<br>ET Epinephrine<br>(n=9) |
| --- | --- | --- | --- | --- |
| ROSC with<br>allocated<br>treatment alone | 1/6*<sup>#</sup> | 9/9 | 0/9*** <sup>##</sup> | 7/9 |
| ROSC in<br>response to<br>rescue IV<br>Epinephrine | 3/5 | not applicable | 6/9 | 1/2 |
| Allocated<br>treatment doses<br>received: |  |  |  |  |
| • 1 dose | 1/6 | 9/9 | 0/9 | 0/9 |
| • 2 doses | 0/6 | 0/9 | 0/9 | 5/9 |
| • 3 doses | 5/6 | 0/9 | 9/9 | 4/9 |
| Rescue IV<br>Epinephrine<br>doses received: |  |  |  |  |
| • 1 dose | 3/6 | 0/9 | 3/9 | 1/9 |
| • 2 doses | 2/6 | 0/9 | 6/9 | 1/9 |
| Mean time (SD)<br>to ROSC,<br>seconds | 586 ± 216<sup>***</sup> | 186 ± 33 | 766 ± 75<sup>*** ###</sup> | 463 ± 117<sup>**</sup> |
| Survival to<br>study end | 4/6 | 9/9 | 3/9<sup>*#</sup> | 8/9 |

ROSC: return of spontaneous circulation. Survival to study end was assessed at 60 minutes
after ROSC. Time to ROSC was measured from initiation of ventilation.
Histological analysis was possible for lambs achieving ROSC only, with availability of tissue
in the following numbers per group: Saline (n=3), IV Epinephrine (n=8), Standard-dose ET
Epinephrine (n=5) or High-dose ET Epinephrine (n=8).
\* p < 0.01 vs IV Epinephrine. \*\* p < 0.001 vs IV Epinephrine. \*\*\* p < 0.0001 vs IV Epinephrine.

### PHYSIOLOGY DURING CPR

The response to treatment of the individual lambs in each group is shown in Figure 2. Diastolic blood pressure significantly increased after allocated treatment in comparison with chest compressions alone in the IV Epinephrine and High-dose ET Epinephrine groups (p <0.0001), but this was not seen in the Saline or Standard-dose ET Epinephrine groups. Diastolic blood pressure significantly increased after rescue IV Epinephrine doses in the Saline group and in both ET Epinephrine groups (p<0.0001).

**Figure 2:**
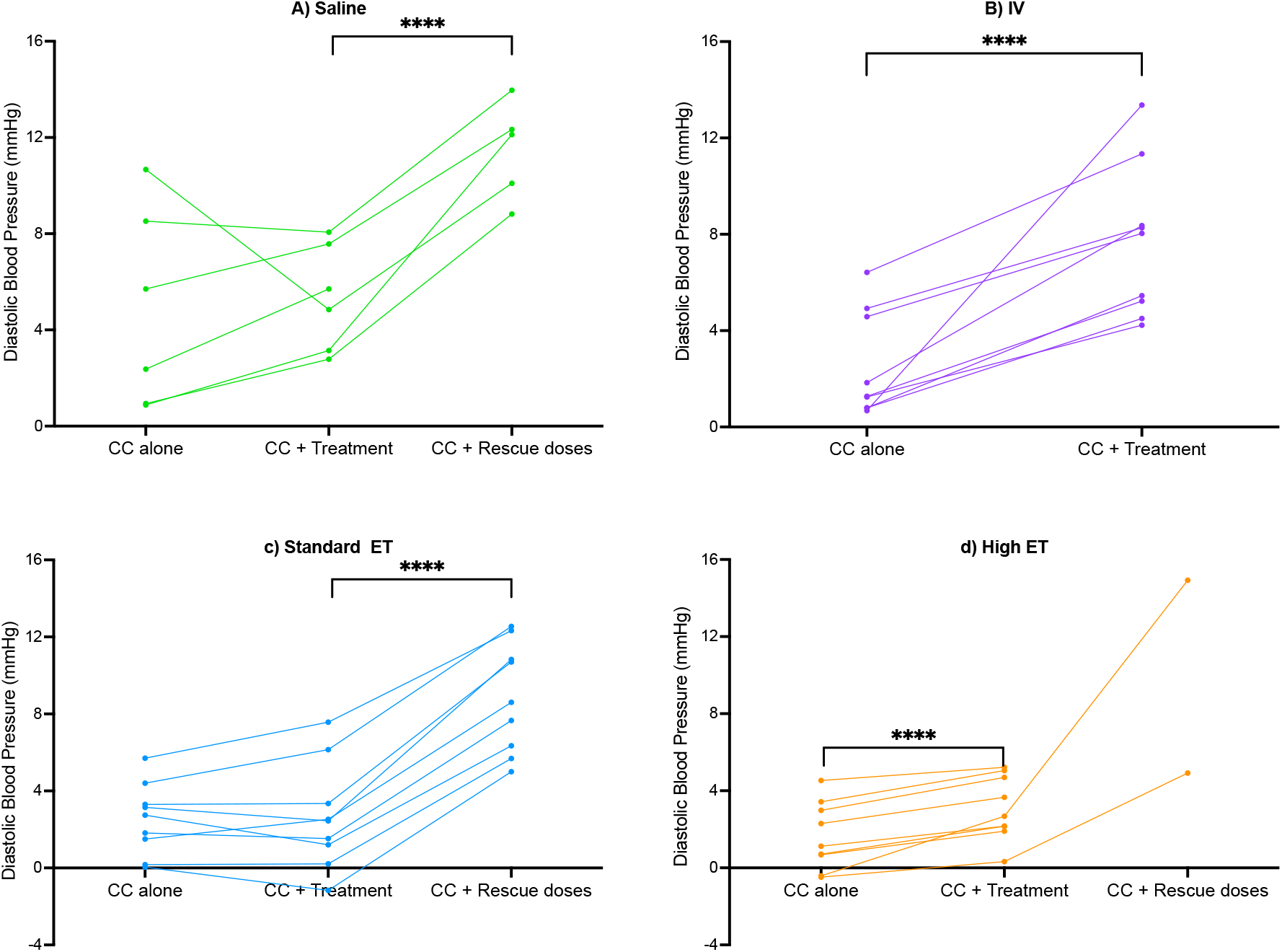
Individual changes to diastolic blood pressure during CPR. Mean diastolic blood pressure of individual lambs during chest compressions (CC) alone, after 1-3 doses of allocated treatment (CC + treatment), and in conjunction with IV rescue epinephrine administration (CC + Rescue doses) in lambs administered A) Saline, B) IV Epinephrine, C) Standard-dose ET Epinephrine and D) High-dose ET Epinephrine. **** indicates p<0.0001. A two-way repeated measures ANOVA with “group” and “time” as the factors was used to compare groups during CPR.

Mean values for mean and diastolic blood pressure, and pulmonary and carotid blood flow, are shown in Figure 3. Changes in pulmonary blood flow during resuscitation were less apparent, with a significant increase after allocated treatment in the IV Epinephrine group (p<0.0001), and after rescue IV Epinephrine was given to the Standard-dose ET Epinephrine group (p<0.05).

**Figure 3:**
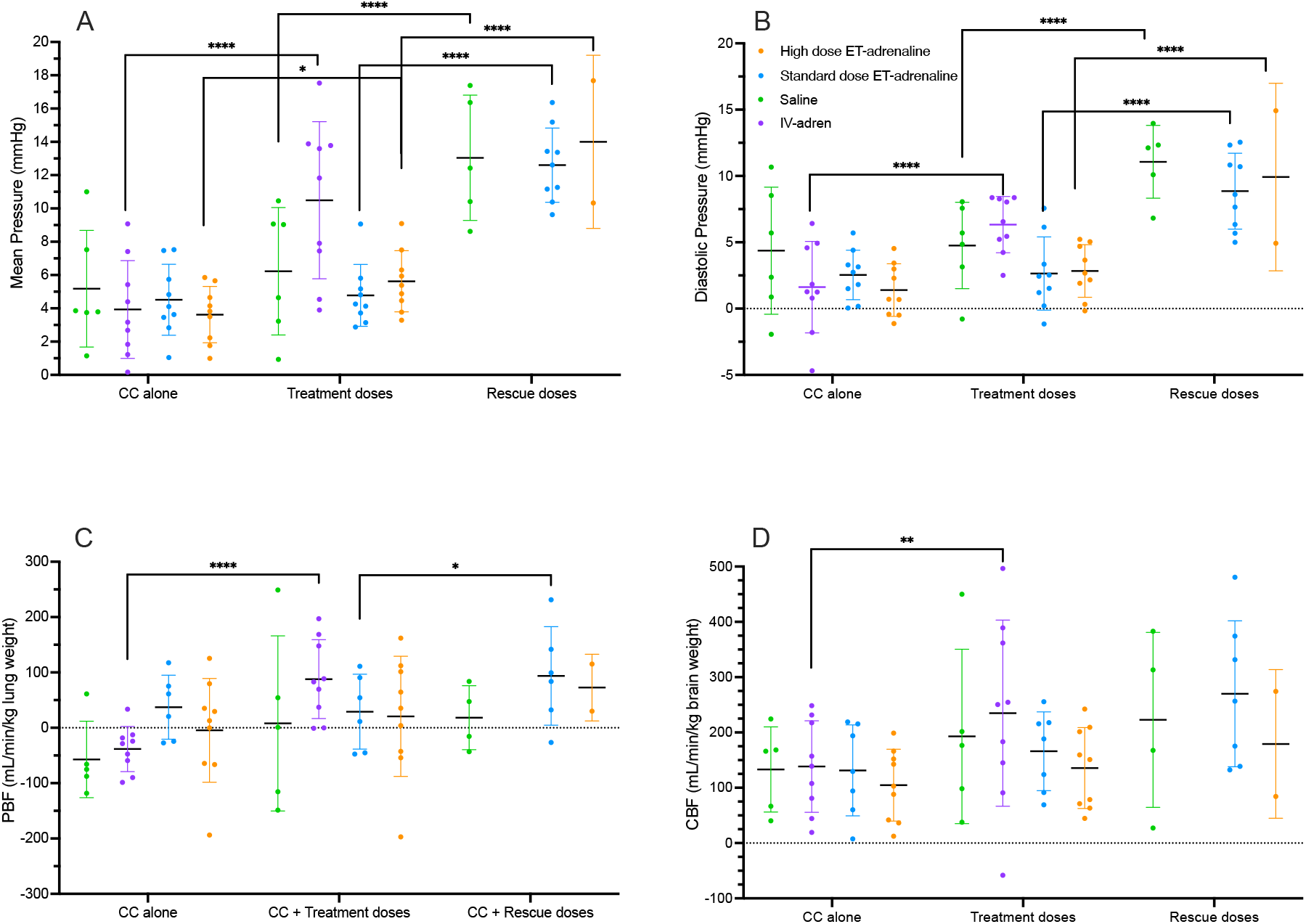
Physiology during CPR. A) mean blood pressure, B) Diastolic blood pressure, C) mean pulmonary blood flow (PBF) and D) mean carotid artery blood flow (CBF) during chest compressions (CC) alone, after allocated treatment (CC + treatment) and in response to rescue IV epinephrine (CC + Rescue doses) in lambs administered Saline (green), IV Epinephrine (purple), Standard-dose ET Epinephrine (blue) and High-dose ET Epinephrine (orange). * indicates p<0.05. ** indicates p<0.01. *** indicates p<0.001. **** indicates p<0.0001. Data are presented as mean ± SD with individual data points for each lamb shown. A two-way repeated measures ANOVA with was used to compare groups during CPR. Note that the comparisons shown are within groups at different time points, while comparisons between groups are not shown.

Similarly, mean carotid blood flow did not change substantially, with a significant increase seen only in the IV Epinephrine group in response to allocated treatment (p<0.01).

### PHYSIOLOGICAL STABILITY AFTER ROSC

The arterial blood pH of the Standard-dose ET Epinephrine group was significantly lower than the other three treatment groups for the first 20 minutes after ROSC, and was lower than the IV Epinephrine and Saline group at 60 min (Figure 4). From 15 minutes after ROSC, the High-dose ET Epinephrine group had a lower pH than the Saline and IV Epinephrine groups. Mean lactate concentration in the IV Epinephrine group was significantly lower than the saline group for the majority of the study, with minimal differences between the other groups.

**Figure 4:**
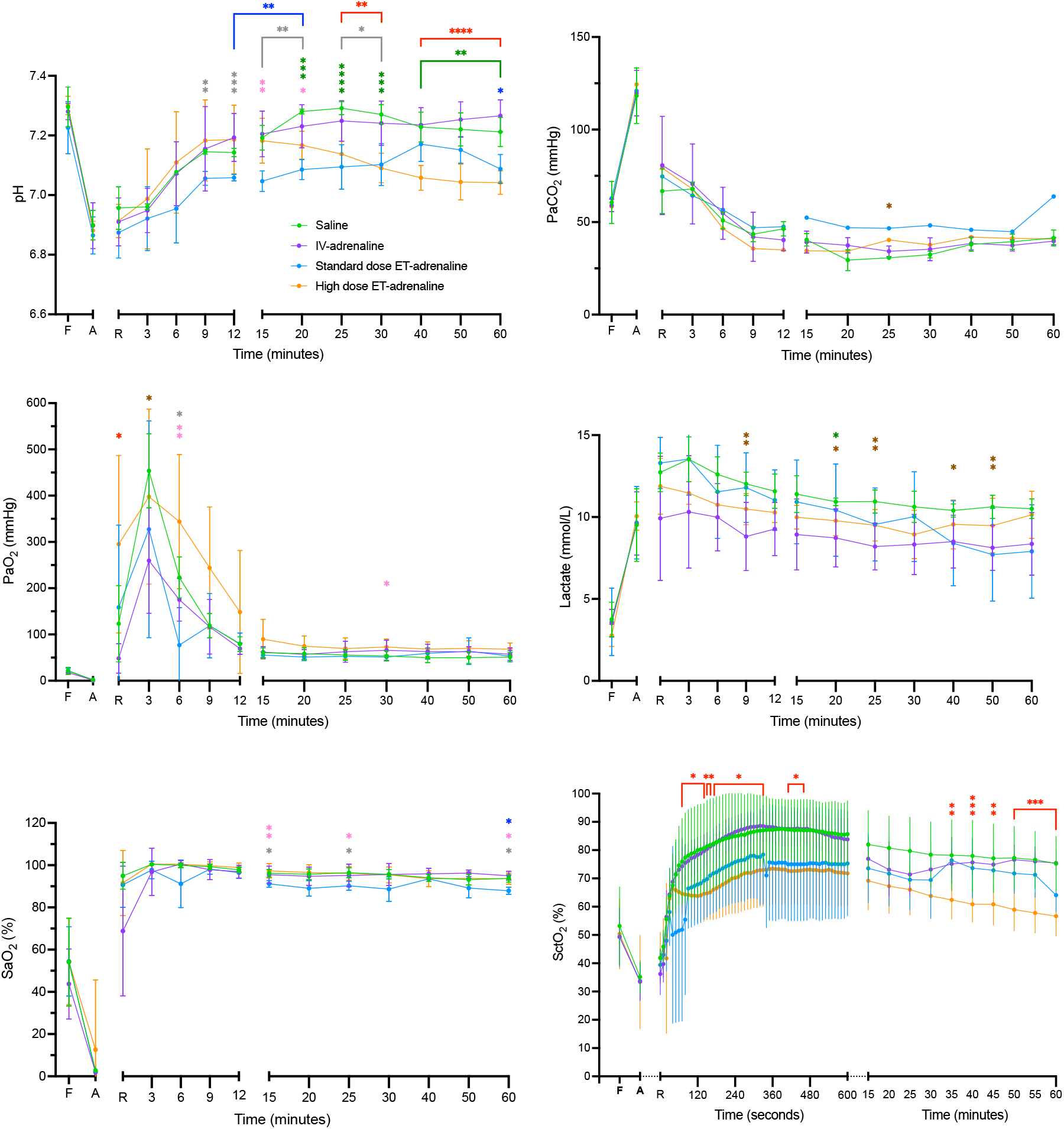
Blood Gas and Oxygenation. A) Arterial pH, B) partial pressure of arterial (Pa) carbon dioxide (PaCO_2_), C) oxygen (PaO_2_), D) arterial lactate concentration, E) arterial oxygen saturation (SaO_2_), F) cerebral oxygenation (SctO_2_) measured at control (fetal, F), end of asphyxia (A), upon ROSC (R) and for one hour of the study. Data are shown for lambs that achieved ROSC: Saline (n=4, green), IV Epinephrine (n=9, purple), Standard-dose ET Epinephrine (n=6, blue) or High-dose ET Epinephrine (n=8, orange). Data are presented as mean ± SD. A two-way repeated measures ANOVA with Holm-Sidak *post hoc* comparison was used to compare post-ROSC physiological data. * indicates p<0.05. ** indicates p<0.01. *** indicates p<0.001. **** indicates p<0.0001. Grey 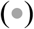 indicates statistical significance between the Saline and Standard-dose ET Epinephrine groups. Dark blue 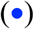 indicates statistical significance between the IV Epinephrine and ET Epinephrine groups. Light pink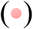 indicates statistical significance between the Standard-dose ET Epinephrine and High-dose ET Epinephrine groups. Dark green 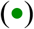 indicates statistical significance between the Saline and High-dose ET Epinephrine groups. Red 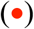 indicates statistical significance between the IV Epinephrine and High-dose ET Epinephrine groups. Brown 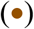 indicates statistical significance between the IV Epinephrine and Saline groups.

The mean partial pressures of oxygen and carbon dioxide in arterial blood, and arterial oxygen saturation levels were similar in all four groups for the majority of the study duration. The mean regional cerebral tissue oxygenation (SctO_2_) was significantly higher in the IV Epinephrine and Saline groups, compared with the ET Epinephrine groups, for the majority of the study duration.

Relative to the IV Epinephrine group, all three other treatment groups were observed to have significantly higher mean blood pressure at multiple time points throughout the study (Figure 5). These increases were most pronounced for the High-dose ET Epinephrine group in the first 25 minutes after ROSC and for the Standard-dose ET Epinephrine group in the final 25 minutes of the study.

**Figure 5:**
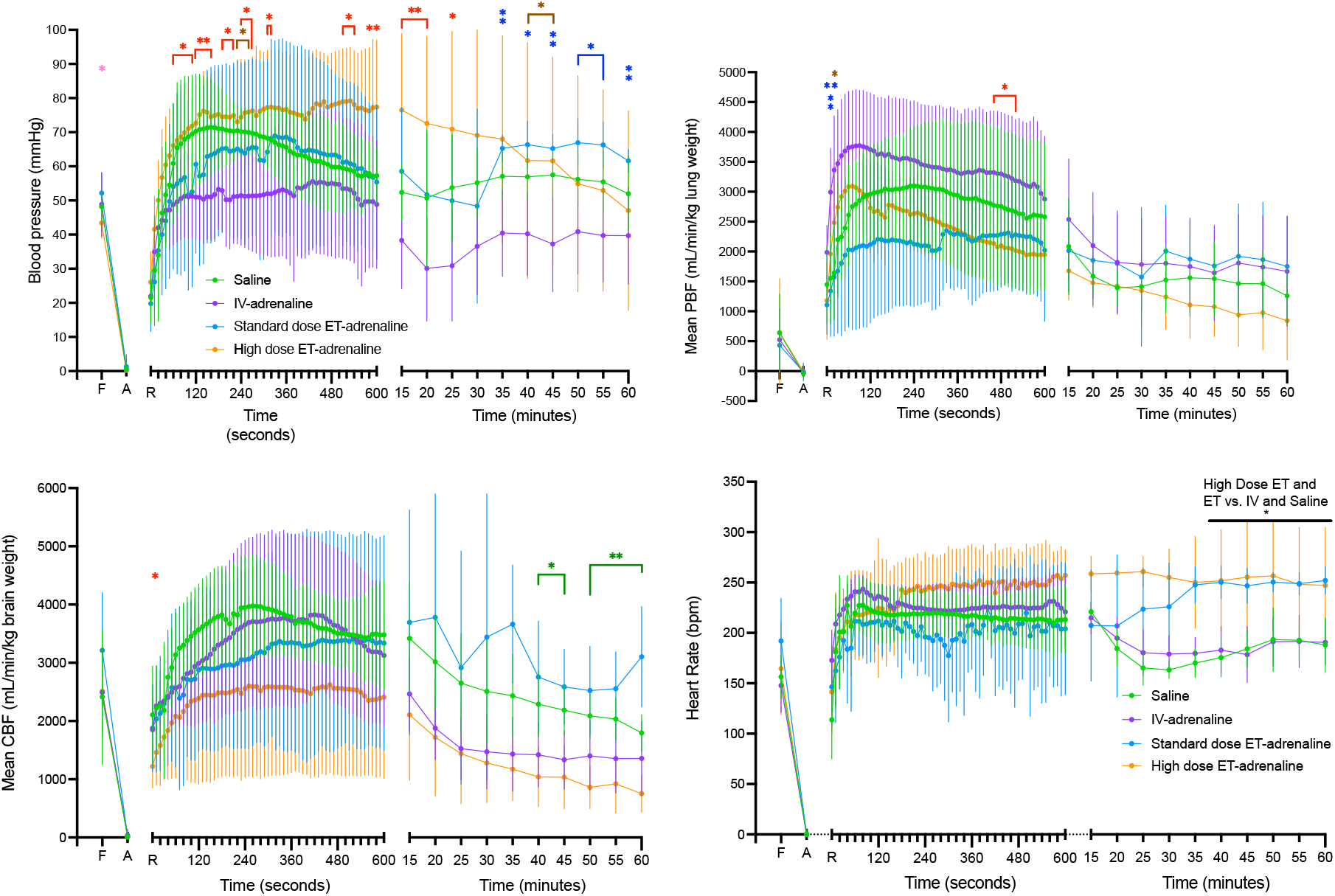
Physiology after ROSC. A) Mean systemic blood pressure, B) mean pulmonary blood flow (PBF) and C) mean carotid arterial blood flow (CBF) and D) Heart rate, measured at control (fetal, F), end of asphyxia (A), upon ROSC (R) and for one hour after ROSC. Data are shown for lambs that achieved ROSC: Saline (n=4, green), IV Epinephrine (n=9, purple), Standard-dose ET Epinephrine (n=6, blue) or High-dose ET Epinephrine (n=8, orange). Data are presented as mean ± SD. * indicates p<0.05. ** indicates p<0.01. Light pink 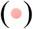 indicates statistical significance between the Standard-dose ET Epinephrine and High-dose ET Epinephrine groups. Red 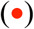 indicates statistical significance between the IV Epinephrine and High-dose ET Epinephrine groups. Brown 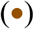 indicates statistical significance between the IV Epinephrine and Saline groups. Dark blue 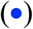 indicates statistical significance between the IV Epinephrine and Standard-dose ET Epinephrine groups.

Mean carotid and pulmonary blood flow were similar in all groups for the majority of the study. Heart rate tended to be higher in the High-dose ET group compared to the others throughout the study (p=0.060) and both ET Epinephrine groups had significantly higher heart rates than the IV Epinephrine and saline groups from 45 minutes.

### MICRO HEMORRHAGE

The occurrence of cerebral micro-hemorrhage in the resuscitated lambs in the four assessed brain regions is expressed as a dichotomous outcome in Table 3, with quantitative findings shown in Supplementary Figure 1. The High-dose ET Epinephrine group had an increased rate of microbleeds relative to the Saline and IV Epinephrine groups. Findings in the other brain regions were similar in all groups.

**Table 3.**
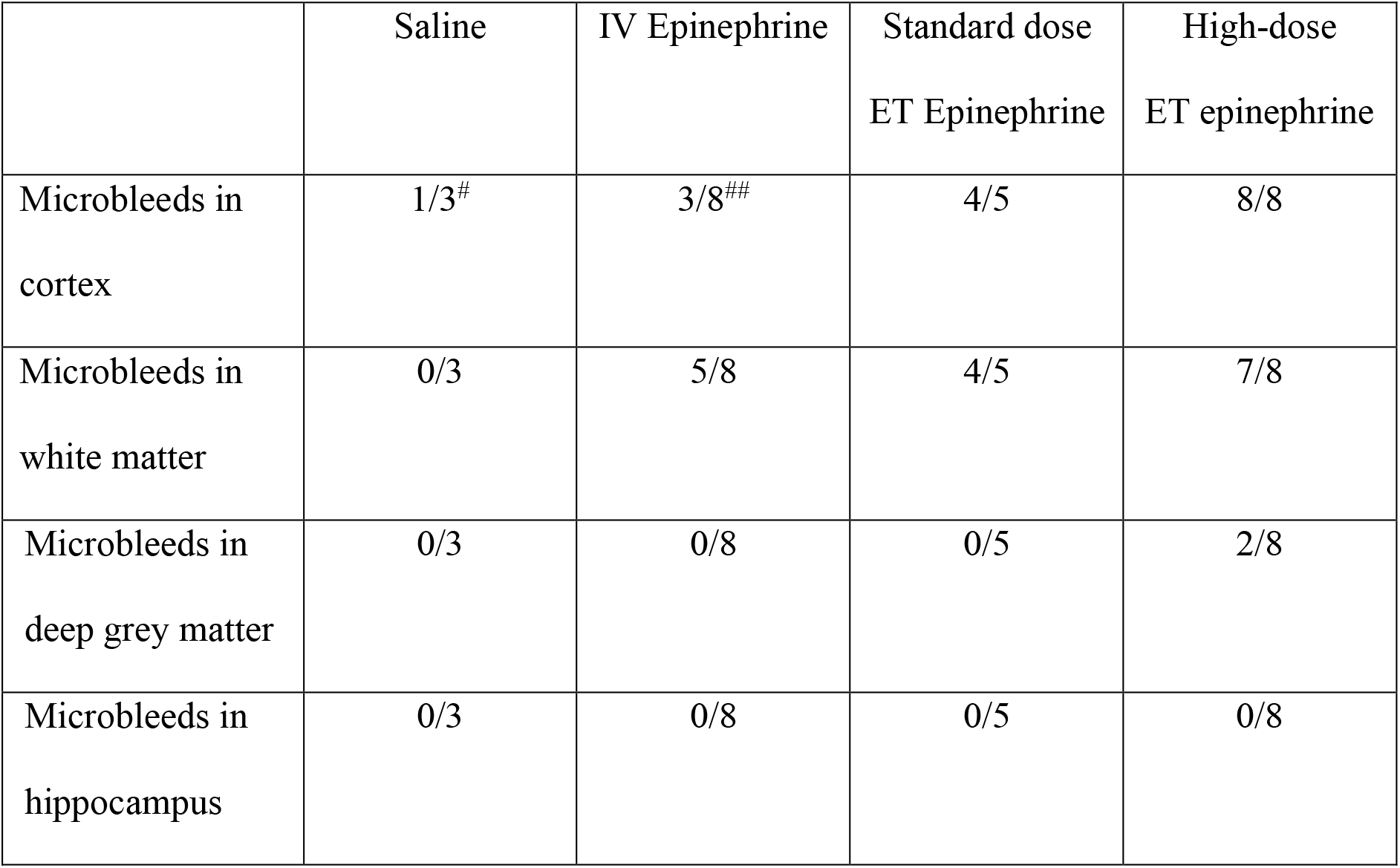
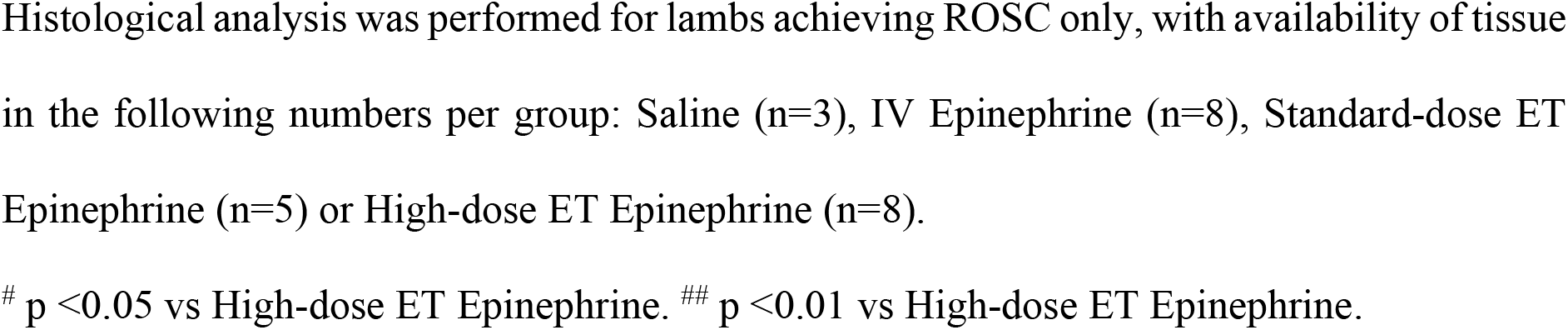
Brain histology and organ weights.

|  | Saline | IV Epinephrine | Standard dose<br>ET Epinephrine | High-dose<br>ET epinephrine |
| --- | --- | --- | --- | --- |
| Microbleeds in<br>cortex | 1/3<sup>#</sup> | 3/8<sup>##</sup> | 4/5 | 8/8 |
| Microbleeds in<br>white matter | 0/3 | 5/8 | 4/5 | 7/8 |
| Microbleeds in<br>deep grey matter | 0/3 | 0/8 | 0/5 | 2/8 |
| Microbleeds in<br>hippocampus | 0/3 | 0/8 | 0/5 | 0/8 |

Histological analysis was performed for lambs achieving ROSC only, with availability of tissue in the following numbers per group: Saline (n=3), IV Epinephrine (n=8), Standard-dose ET Epinephrine (n=5) or High-dose ET Epinephrine (n=8).

### PLASMA EPINEPHRINE CONCENTRATION

Plasma epinephrine levels were measured in the two ET Epinephrine groups. These are shown in Supplementary Figure 2. High-dose ET Epinephrine lambs had significantly higher plasma epinephrine levels from 3 minutes compared to the Standard dose ET lambs. Standard dose ET Epinephrine lambs showed an increase to mean (SD) 45 ± 20 ng/ml at ROSC, which gradually decreased over 15 minutes. In contrast, the level in High-dose ET Epinephrine lambs was substantially higher, reaching 157 ± 120 ng/ml at 3 minutes and remaining elevated at >150 ng/ml for at least 15 minutes, at which time it was 178 ± 118 ng/ml.

## Discussion

We conducted this study to evaluate the effectiveness of ET epinephrine in achieving ROSC and maintaining cardiovascular stability after ROSC, in asystolic asphyxiated newborn lambs. Our findings indicate that administration of ET epinephrine at the current maximum recommended “standard dose” (100 micrograms/kg) is ineffective, with none of the lambs allocated to this treatment group achieving ROSC, without the additional use of ‘rescue’ IV epinephrine. These outcomes contrasted with the lambs allocated to initial IV Epinephrine treatment, where ROSC was achieved with allocated treatment in all cases. In particular, lambs receiving Standard-dose ET epinephrine showed no improvement compared with the Saline placebo group, where 1/6 lambs achieved ROSC without rescue IV treatment.

It is likely that the poor response in the Standard-dose ET Epinephrine treated lambs is a consequence of limited bioavailability of the administered epinephrine to the myocardium. This could be due to the presence of airway liquid and low pulmonary blood flow, which is a common feature of newborns as they transition from fetal to newborn life.(11, 12) This concept is supported by the observation that during CPR, the mean and diastolic blood pressures observed in Standard-dose ET Epinephrine treated lambs during chest compressions did not increase, relative to the values seen during chest compressions alone. This, again, was similar to the findings in the Saline placebo group.

In contrast to our initial hypothesis, the use of High-dose ET Epinephrine resulted in a rate of ROSC following allocated treatment (7/9) that was significantly greater than that obtained with Standard-dose ET Epinephrine or Saline, and close to that obtained with IV Epinephrine. This suggests that increasing the administered dose by a magnitude of ten is sufficient to overcome some of the limitations in absorption from the airways into the central circulation, and produce the desired physiological effect on the heart. This was apparent in the physiological observations during CPR, where the diastolic blood pressure significantly increased during the period of chest compressions and allocated treatment, in comparison with the values during chest compressions alone.

Differences were also apparent in the time taken to achieve ROSC. Although slower than the response seen to IV Epinephrine, the High-dose ET Epinephrine group achieved ROSC, on average, approximately 5 minutes earlier than the Standard-dose ET Epinephrine group. In the clinical setting, a reduction in time of asystole by 5 minutes would have the potential to reduce the severity of hypoxic-ischemic brain injury.

Our assessment of plasma epinephrine levels demonstrated that, at the time of ROSC, the level in the High-dose ET Epinephrine group was approximately twice that seen with the standard dose, supporting the concept that greater systemic absorption occurs with elevated ET dosing. We found that both ET groups had elevated plasma epinephrine levels for a sustained period after ROSC, which differs from IV administration, which we have previously observed to peak between 10-20 ng/mL and return to <10 ng/mL by 15 minutes after ROSC, in asphyxiated lambs receiving an IV dose of approximately 10 micrograms/kg body weight.(14, 21) It appears that much of the ET Epinephrine dose administered remains in the airways during CPR, but is absorbed after ROSC is achieved, producing persistently high plasma levels. Whilst the Standard-dose ET Epinephrine group had levels that were clearly decreasing by 15 minutes after ROSC, the level seen in High-dose ET Epinephrine lambs did not show any apparent decrease between 3 and 15 minutes after ROSC. Sustained exposure to such high epinephrine levels could have adverse effects on the heart, and other vital organs, in the post-resuscitation period.

Some of the other physiological variables we evaluated suggested that the lambs exposed to ET Epinephrine were not recovering as effectively after resuscitation. Both ET Epinephrine groups demonstrated a lower pH relative to the IV Epinephrine group. Both ET Epinephrine groups had a much higher blood pressure after ROSC, and a lower SctO_2_. These findings could be explained by a sustained vasoconstriction effect due to epinephrine, which could adversely impact tissue perfusion, most concerningly to the brain. The elevation in blood pressure, the prolonged period of asphyxia before ROSC, or both factors in combination, would present a potential explanation for the increased frequency of brain micro-hemorrhage observed in the High-dose ET Epinephrine lambs on histological analysis. Micro-hemorrhage was also seen in all but one of the standard dose ET Epinephrine group that underwent histological examination. Within the Standard-dose ET Epinephrine group, 3/6 lambs that initially achieved ROSC (after rescue IV Epinephrine) subsequently had a second cardiac arrest, resulting in death prior to study endpoint. While the reason for such a deterioration cannot be accurately determined, this group had the longest time to ROSC, mean 766 seconds, and therefore were exposed to the most prolonged period of myocardial ischemia. It is also possible that the elevated plasma epinephrine levels seen after ET Epinephrine administration have an adverse effect on cardiac function and circulatory stability, perhaps influenced by the persistent tachycardia seen in this group. However, the High-dose ET Epinephrine lambs were exposed to substantially higher plasma levels, had a similar degree of tachycardia, and all 8 lambs achieving ROSC survived for 60 minutes. It may be that elevated epinephrine levels are more likely to produce adverse effects in the context of greater ischaemic damage to the myocardium.

While there are limitations in how preclinical data can be applied to the clinical setting, and it is unclear how some of the outcomes we identified would translate into clinical outcomes in newborn infants (e.g., the impact of micro-hemorrhage on neurodevelopment), this study has several strengths, including the use of a well-established protocol for asphyxia and resuscitation in ovine studies, and the randomised allocation to the four treatment groups.

Other preclinical studies of ET Epinephrine have similarly shown reduced rates of ROSC, when used at the current maximum recommended “standard dose” (100 micrograms/kg) without the use of rescue IV Epinephrine. Songstad reported a rate of ROSC in response to allocated treatment of 2/5 with “standard dose” ET Epinephrine versus 6/6 with IV Epinephrine.(14) Vali treated a total of 22 lambs with ET Epinephrine, either before or after initiating CPR, and 12/22 achieved ROSC without IV rescue dosing, whereas lambs receiving IV Epinephrine (either via UVC or RA) achieved ROSC in 19/22 cases.(13) While the exact rates of ROSC observed differ slightly between studies, which may be accounted for by slight variations in treatment protocols or study population, a clear pattern is evident that ET Epinephrine administration is less likely to achieve ROSC than IV Epinephrine treatment.

ET Epinephrine administration was included in the first published version of consensus resuscitation recommendations specific to the newborn infant, produced by ILCOR in 1999.(22) At this time, the recommended dose for either the ET or IV group was identical (10-30 micrograms/kg), and neither route was specifically identified as preferred, although the recommendation document did highlight “concerns that the endotracheal route may not result in as effective a level of epinephrine as does the intravenous route”. On revision of these recommendations in 2005,(23) the authors noted “past guidelines recommended that initial doses of epinephrine be given through an endotracheal tube because the dose can be administered more quickly”, but highlighted that the only preclinical study using a dose of 10-30 micrograms/kg, conducted in piglets at 2-4 days of age, showed no effect.(24) They therefore recommended that IV was the preferred route for epinephrine administration, and advised that while IV access was being obtained, a higher dose (up to 100 micrograms/kg) could be considered.(23) Updates to these guidelines published between 2010 and 2020 (the most recent version) have consistently identified IV administration as the preferred route, but included ET Epinephrine as an option if intravenous access is not yet available.(4, 25) The 2020 document specifically highlights that “administration of endotracheal epinephrine (adrenaline) should not delay attempts to establish vascular access”.(4) It remains the case that there are no randomised trial data to support ET Epinephrine use in newborn infants.(15)

Intraosseous access is another option for epinephrine administration. While the 2005 ILCOR document simply stated that “intraosseous lines are not commonly used in newly born infants”, the intraosseous (IO) route has been increasingly acknowledged in more recent revisions. In the 2020 treatment recommendations state that “if umbilical venous access is not feasible, the intraosseous route is a reasonable alternative” and that either the IV or IO route is suitable for neonatal resuscitation outside the delivery room setting.(4) However, the published clinical data for IO use in neonates are restricted to case series and case reports, and important complications have been reported.(26) Subgroup analysis of a recent adult RCT showed that IO and IV Epinephrine produced almost identical rates of ROSC in the setting of out-of- hospital cardiac arrest.(27) A recent preclinical study conducted in asystolic newborn lambs found that IO administration of epinephrine produced similar rates of ROSC (7/9 IO versus 10/12 IV), a similar time taken to achieve ROSC, and similar plasma epinephrine levels.(21) Our findings raise questions about the role of ET Epinephrine in newborn resuscitation. The efficacy of the ET route has been consistently lower in achieving ROSC than that obtained with IV Epinephrine, in this study, and other recent preclinical studies of asystolic lambs.(13, 14) Published clinical neonatal data do not include any randomised trials. Observational data, despite including limited numbers of infants, have consistently shown that a minority of infants improve after ET Epinephrine, with high rates of additional IV Epinephrine administration prior to achievement of ROSC.(15-17) Whilst there is a long-established historical precedent for ET Epinephrine use in neonatal resuscitation, it is unlikely that any new treatment would be readily accepted into treatment guidelines if it were associated with a similar profile of clinical and preclinical outcomes. Neonatal resuscitation guidelines currently include two alternatives, the IV route, which is clearly preferrable, and IO access, which despite being limited to clinical case series evidence in neonates, has similar preclinical outcomes to IV treatment, and is widely used in resuscitation of older patients.(4, 21) Given this context, should ET Epinephrine, at the current dose, continue to feature in consensus guidance?

In contrast, our finding that a much higher dose of ET Epinephrine, 1 mg/kg, can produce similar rates of ROSC to IV Epinephrine treatment may have some promise. However, it is unclear whether this approach is ready for translation into a clinical trial. Although 7/9 High-dose ET Epinephrine lambs achieved ROSC without rescue IV Epinephrine, all required either two or three doses to do so. The need for multiple doses may negate any advantage in time to administration that the ET route presents over other alternatives.(6)

The evidence of micro-hemorrhage associated with High-dose Epinephrine in the preclinical setting also raises the concern of adverse neurological effects in those that achieve ROSC. A trial of IV Epinephrine versus placebo for adult out-of-hospital cardiac arrest identified such a ‘trade-off’ situation, as epinephrine use significantly increased survival, but also significantly increased severe neurological impairment in survivors.(28)

Given the urgency, and infrequent and unpredictable nature of epinephrine use at birth, clinical trials may be better focused on comparing routes with higher likelihood of achieving ROSC, such as IV and IO administration. Further preclinical studies, focused on dose-finding, or evaluation of epinephrine formulations with enhanced absorption characteristics, may identify an approach to ET epinephrine that has more promise for clinical assessment.

## Conclusions

In this preclinical randomised study assessing different routes of epinephrine administration in asystolic newborn lambs, we found the current recommended “standard dose” of ET Epinephrine to be significantly less effective in achieving ROSC than IV Epinephrine, performing similarly to saline placebo. This finding, in the context of other preclinical data and observational clinical data showing reduced efficacy, calls into question whether ET Epinephrine administration should be recommended in neonatal resuscitation guidelines. The use of a higher ET dose, 1 mg/kg epinephrine, produced a high rate of ROSC similar to that seen with IV administration, but with evidence of increased micro-haemorrhage in survivors. Generation of high quality preclinical and clinical data to guide the use of epinephrine during neonatal resuscitation is a priority.

**What is known?**

- ILCOR neonatal resuscitation recommendations suggest administering Endotracheal Epinephrine for infants with heart rate of 60 beats per minute or less, if intravascular access is not yet available
- The suggested Endotracheal Epinephrine dose is higher than the standard IV Epinephrine dose (50-100 micrograms/kg, versus 10-20 micrograms/kg) due to concerns about absorption into the systemic circulation
- There are no randomized clinical trial data to support Endotracheal Epinephrine use in newborns, and observational studies suggest most infants subsequently also require IV Epinephrine before achieving return of spontaneous circulation (ROSC)

**What new information does this article contribute?**

- Endotracheal epinephrine, administered at the standard dose currently recommended in neonatal resuscitation guidelines (100 micrograms/kg), was ineffective in achieving ROSC in a preclinical study of asystolic newborn lambs (0/9 responded to allocated treatment)
- High-dose Endotracheal Epinephrine (1 mg/kg) produced similar rates of ROSC to IV Epinephrine (7/9 versus 9/9 responded to allocated treatment)
- Endotracheal Epinephrine-treated lambs had lower pH and cerebral oxygen saturations, and elevated blood pressure and lactate levels after ROSC, and High-dose Endotracheal Epinephrine resulted in an increased rate of micro-hemorrhage

## Novelty and Significance

In this randomized preclinical study, we found that the currently recommended ‘Standard-dose’ (100 micrograms/kg) of Endotracheal Epinephrine was ineffective in achieving ROSC in asystolic newborn lambs, performing similarly to the use of IV saline placebo. Lambs receiving Standard-dose Endotracheal Epinephrine achieved ROSC only after rescue IV Epinephrine use. We evaluated a novel High-dose of Endotracheal Epinephrine (1 mg/kg), finding that it produced rates of ROSC similar to IV Epinephrine, indicating that increasing the administered dose may overcome issues with systemic absorption in the fluid-filled newborn lung. However, both Endotracheal Epinephrine groups demonstrated adverse effects after ROSC, including low pH and cerebral oxygen saturations, and elevated lactate and blood pressure. Given the lack of clinical data suggesting efficacy, and the challenges of conducting clinical trials in this population of infants, these findings from a randomized preclinical study call into question whether Standard-dose Endotracheal Epinephrine should continue to be recommended in neonatal resuscitation guidelines. Use of a higher Endotracheal Epinephrine dose, while potentially promising in achieving ROSC, requires further evaluation of both efficacy and post-resuscitation effects before any consideration of clinical translation.

## ACKNOWLEDGEMENTS

The authors would like to thank Alison Moxham, Valerie Zahra and Karyn Rodgers for their technical support.

## CONTRIBUTOR STATEMENT

All named authors contributed to one or more of: conception and design of the study, data acquisition, analysis and interpretation of the data. GRP, YB and CTR co-wrote the first draft of the manuscript. All authors revised the final manuscript and approved it prior to submission. The authors have no conflicts of interest to disclose.

## FUNDING

This research was supported by National Health and Medical Research Council (NHMRC) Project Grant APP1158494 and Fellowships (GRP: APP1173731, SBH:APP545921, SM:APP1136216, CTR:APP1175634), a National Heart Foundation of Australia Vanguard Grant (103022) and the Victorian Government’s Operational Infrastructure Support Program.

## PROTOCOL AND DATA ACCESS

A protocol was included in research ethics submission, but was not publicly registered. Data access will be considered on reasonable request to the authors.

## SUPPLEMENTARY FIGURES (for online publication)

**Supplementary Figure 1:**
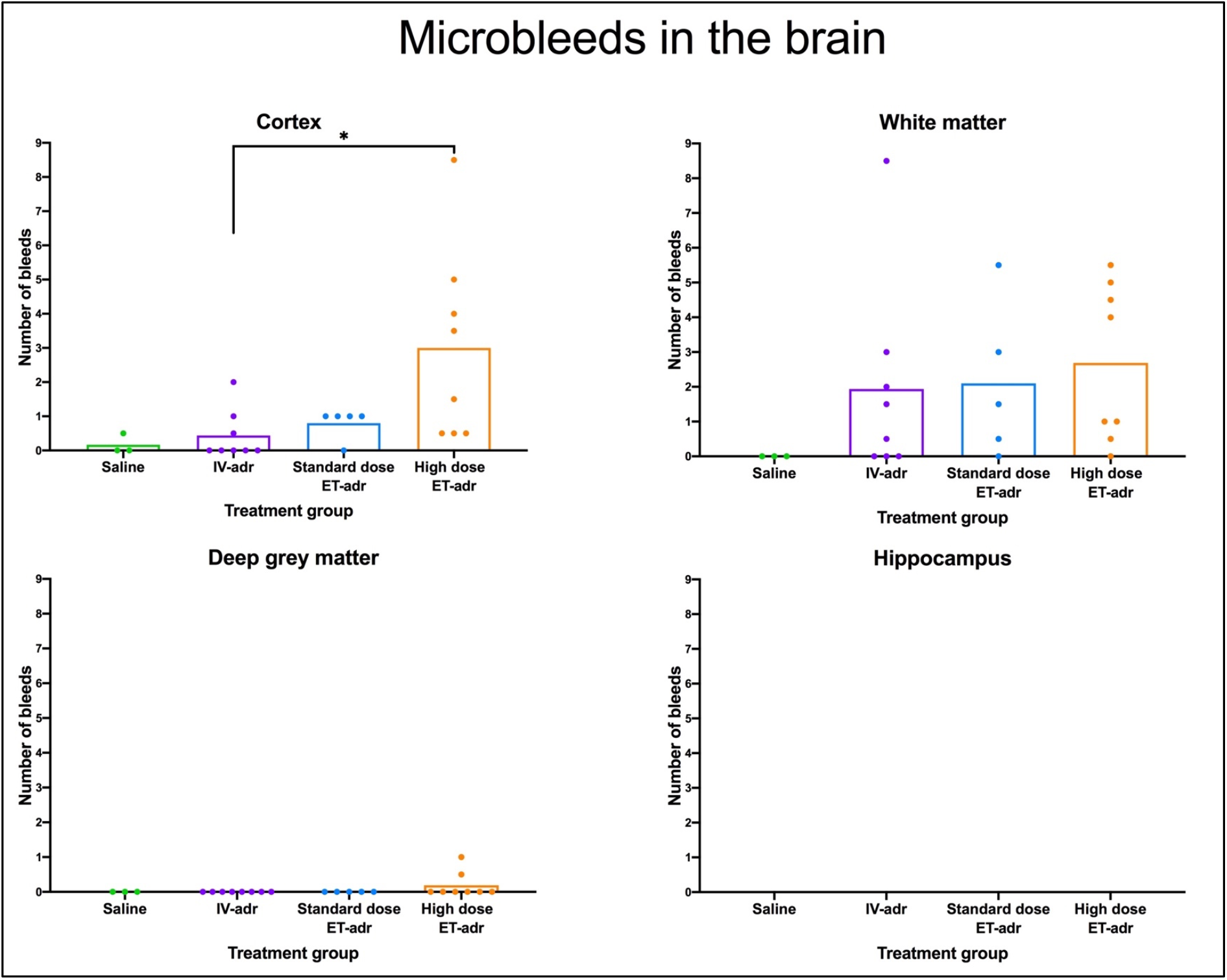
Number of microbleeds in different brain regions the resuscitated lambs. Data are shown for lambs which achieved ROSC: saline (n=3), IV Epinephrine (n=8), Standard-dose ET Epinephrine (n=5) or High-dose ET Epinephrine (n=8). The number of bleeds in each animal is an average of the number of bleeds found on duplicated slides. Data are presented as mean. Significance was measured by multiple comparisons one-way ANOVA. * indicates p<0.05. *IV: intravenous; ET: endotracheal; Adr: adrenaline*

**Supplementary Figure 2:**
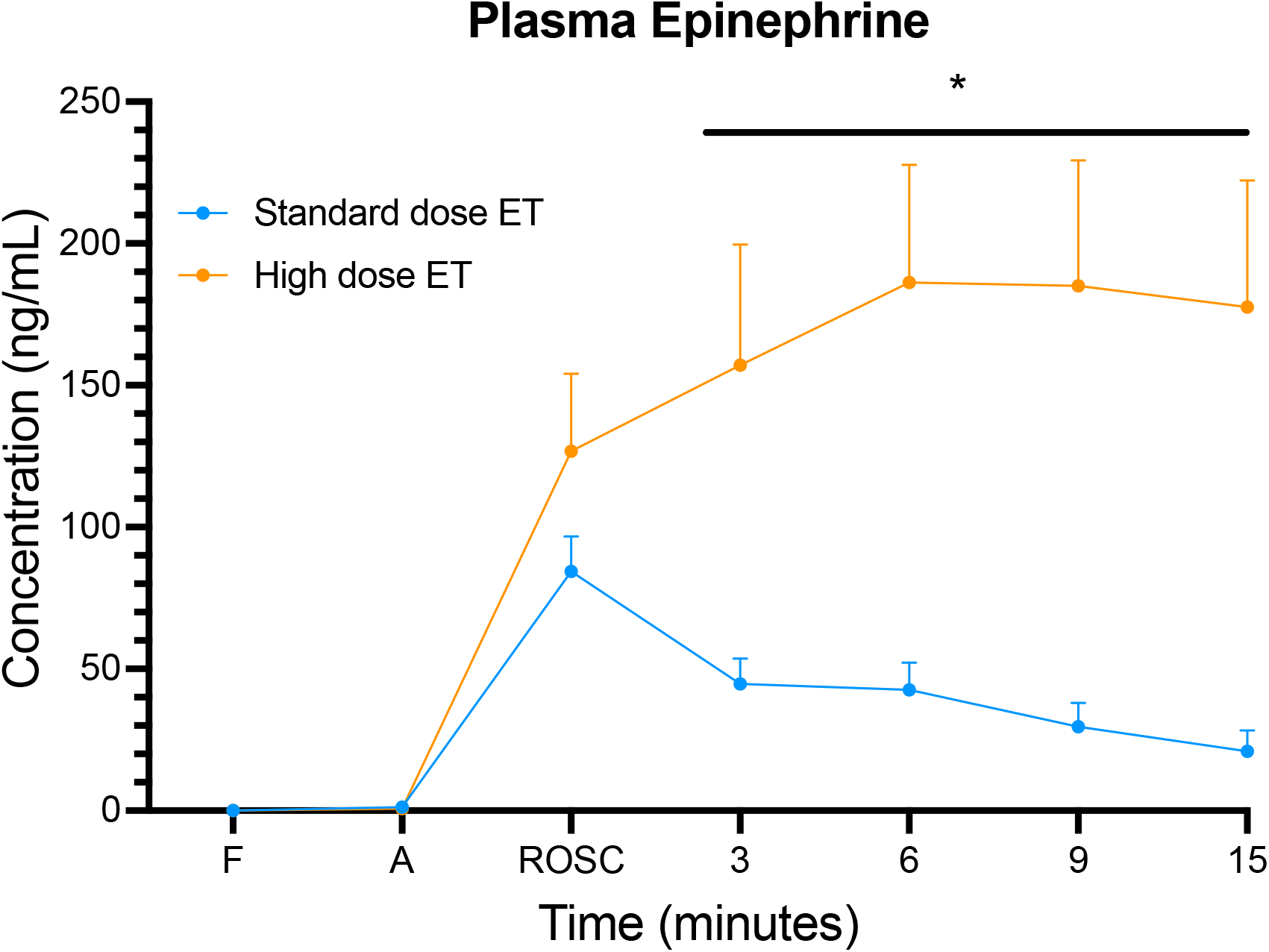
Plasma epinephrine concentration of asphyxiated lambs. Data are shown for lambs which achieved ROSC: Standard-dose ET Epinephrine (n=5), High-dose ET Epinephrine (n=8). Data are presented as mean ± SD. A two-way repeated measures ANOVA with Holm-Sidak *post hoc* comparison was used to compare the plasma epinephrine levels. * indicates p<0.05. *F: fetal; A: end of asphyxiation; ROSC: return of spontaneous circulation*.

## Notes

### Competing Interest Statement

The authors have declared no competing interest.

## References

1. Lawn JE, Blencowe H, Oza S, You D, Lee ACC, Waiswa P, et al. Every Newborn: progress, priorities, and potential beyond survival. The Lancet. 2014;384(9938):189–205.

2. Lee AC, Kozuki N, Blencowe H, Vos T, Bahalim A, Darmstadt GL, et al. Intrapartum-related neonatal encephalopathy incidence and impairment at regional and global levels for 2010 with trends from 1990. Pediatric research. 2013;74 Suppl 1:50–72.

3. O’Donnell AI, Gray PH, Rogers YM. Mortality and neurodevelopmental outcome for infants receiving adrenaline in neonatal resuscitation. Journal of paediatrics and child health. 1998;34(6):551–6.

4. Wyckoff MH, Wyllie J, Aziz K, de Almeida MF, Fabres J, Fawke J, et al. Neonatal Life Support: 2020 International Consensus on Cardiopulmonary Resuscitation and Emergency Cardiovascular Care Science With Treatment Recommendations. Circulation. 2020;142(16_suppl_1):S185–S221.

5. Madar J, Roehr CC, Ainsworth S, Ersdal H, Morley C, Rudiger M, et al. European Resuscitation Council Guidelines 2021: Newborn resuscitation and support of transition of infants at birth. Resuscitation. 2021;161:291–326.

6. McKinsey S, Perlman JM. Resuscitative interventions during simulated asystole deviate from the recommended timeline. Archives of disease in childhood Fetal and eonatal edition. 2016;101(3):F244–7.

7. Redding JS, Asuncion JS, Pearson JW. Effective routes of drug administration during cardiac arrest. Anesthesia and analgesia. 1967;46(2):253–8.

8. Roberts JR, Greenberg MI, Knaub MA, Kendrick ZV, Baskin SI. Blood levels following intravenous and endotracheal epinephrine administration. JACEP. 1979;8(2):53–6.

9. Crespo SG, Schoffstall JM, Fuhs LR, Spivey WH. Comparison of two doses of endotracheal epinephrine in a cardiac arrest model. Ann Emerg Med. 1991;20(3):230–4.

10. Roberts JR, Greenburg MI, Knaub M, Baskin SI. Comparison of the pharmacological effects of epinephrine administered by the intravenous and endotracheal routes. JACEP. 1978;7(7):260–4.

11. Hooper SB, Harding R. Role of aeration in the physiological adaptation of the lung to air-breathing at birth. Current Respiratory Medicine Reviews. 2005;1:185–95.

12. Polglase GR, Hooper SB. Role of Intra-luminal Pressure in Regulating PBF in the Fetus and After Birth. Current Pediatric Reviews. 2006;2(4):287–99.

13. Vali P, Chandrasekharan P, Rawat M, Gugino S, Koenigsknecht C, Helman J, et al. Evaluation of Timing and Route of Epinephrine in a Neonatal Model of Asphyxial Arrest. J Am Heart Assoc. 2017;6(2).

14. Songstad NT, Klingenberg C, McGillick EV, Polglase GR, Zahra V, Schmolzer GM, et al. Efficacy of Intravenous, Endotracheal, or Nasal Adrenaline Administration During Resuscitation of Near-Term Asphyxiated Lambs. Front Pediatr. 2020;8:262.

15. Isayama T, Mildenhall L, Schmölzer GM, Kim H-S, Rabi Y, Ziegler C, et al. The Route, Dose, and Interval of Epinephrine for Neonatal Resuscitation: A Systematic Review. Pediatrics. 2020;146(4).

16. Barber CA, Wyckoff MH. Use and efficacy of endotracheal versus intravenous epinephrine during neonatal cardiopulmonary resuscitation in the delivery room. Pediatrics. 2006;118(3):1028–34.

17. Halling C, Sparks JE, Christie L, Wyckoff MH. Efficacy of Intravenous and Endotracheal Epinephrine during Neonatal Cardiopulmonary Resuscitation in the Delivery Room. The Journal of pediatrics. 2017;185:232–6.

18. Kilkenny C, Browne WJ, Cuthill IC, Emerson M, Altman DG. Improving bioscience research reporting: the ARRIVE guidelines for reporting animal research. PLoS Biol. 2010;8(6):e1000412. doi: 10.1371/journal.pbio.

19. Polglase GR, Schmolzer GM, Roberts CT, Blank DA, Badurdeen S, Crossley KJ, et al. Cardiopulmonary Resuscitation of Asystolic Newborn Lambs Prior to Umbilical Cord Clamping; the Timing of Cord Clamping Matters! Front Physiol. 2020;11:902.

20. Craggs LJ, Yamamoto Y, Deramecourt V, Kalaria RN. Microvascular pathology and morphometrics of sporadic and hereditary small vessel diseases of the brain. Brain Pathol. 2014;24(5):495–509.

21. Roberts CT, Klink S, Schmölzer GM, Blank DA, Badurdeen S, Crossley KJ, et al. Comparison of intraosseous and intravenous epinephrine administration during resuscitation of asphyxiated newborn lambs. Archives of Disease in Childhood - Fetal and Neonatal Edition. 2022;107(3):311–6.

22. Kattwinkel J, Niermeyer S, Nadkarni V, Tibballs J, Phillips B, Zideman D, et al. ILCOR advisory statement: resuscitation of the newly born infant. An advisory statement from the pediatric working group of the International Liaison Committee on Resuscitation. Circulation. 1999;99(14):1927–38.

23. Association AH. 2005 American Heart Association Guidelines for Cardiopulmonary Resuscitation and Emergency Cardiovascular Care. Part 13: Neonatal Resuscitation Guidelines. Circulation. 2005;112(24 Suppl):IV188–95.

24. Kleinman ME, Oh W, Stonestreet BS. Comparison of intravenous and endotracheal epinephrine during cardiopulmonary resuscitation in newborn piglets. Critical care medicine. 1999;27(12):2748–54.

25. Perlman JM, Wyllie J, Kattwinkel J, Atkins DL, Chameides L, Goldsmith JP, et al. Part 11: Neonatal resuscitation: 2010 International Consensus on Cardiopulmonary Resuscitation and Emergency Cardiovascular Care Science With Treatment Recommendations. Circulation. 2010;122(16 Suppl 2):S516–38.

26. Scrivens A, Reynolds PR, Emery FE, Roberts CT, Polglase GR, Hooper SB, et al. Use of Intraosseous Needles in Neonates: A Systematic Review. Neonatology. 2019;116(4):305–14.

27. Nolan JP, Deakin CD, Ji C, Gates S, Rosser A, Lall R, et al. Intraosseous versus intravenous administration of adrenaline in patients with out-of-hospital cardiac arrest: a secondary analysis of the PARAMEDIC2 placebo-controlled trial. Intensive care medicine. 2020;46(5):954–62.

28. Perkins GD, Ji C, Deakin CD, Quinn T, Nolan JP, Scomparin C, et al. A Randomized Trial of Epinephrine in Out-of-Hospital Cardiac Arrest. The New England journal of medicine. 2018;379(8):711–21.

